# Ctenophores and parahoxozoans independently evolved complex and functionally diverse sets of voltage-gated K^+^ channels

**DOI:** 10.1101/2024.12.09.627618

**Authors:** Benjamin T. Simonson, Zhaoyang Jiang, Joseph F. Ryan, Timothy Jegla

**Affiliations:** Department of Biology and Huck Institutes of the Life Sciences, Penn State University, University Park, PA; Whitney Laboratory for Marine Bioscience, University of Florida, St. Augustine, FL; Department of Biology, University of Florida, Gainesville, FL

**Author notes:** Authors contributed equally (Benjamin T. Simonson, Zhaoyang Jiang). Correspondence: Timothy Jegla, Department of Biology, Penn State University, University Park, PA.

**Keywords:** Shaker, Ether-a-go-go, KCNQ, ctenophore, K^+^ channel, HCN

## Abstract

The ctenophore species *Mnemiopsis leidyi* is known to have a large set of voltage-gated K^+^ channels, but little is known about the functional diversity of these channels or the evolutionary history of this gene family in other ctenophore species. Here, we searched the genomes of two additional ctenophore species, *Beroe ovata* and *Hormiphora californensis*, for voltage-gated K^+^ channels and functionally expressed a subset of *M. leidyi* channels. We found that the last common ancestor of these three disparate ctenophore lineages probably had at least 33 voltage-gated K^+^ channels. Two of these genes belong to the EAG family, and the remaining 31 belong to the Shaker family and form a single clade within the animal/choanoflagellate Shaker phylogeny. We additionally found evidence for 10 of these Shaker channels in a transcriptome of the early branching ctenophore lineage *Euplokamis dunlapae*, suggesting that the diversification of these channels was already underway early in ctenophore evolution. We functionally expressed 16 *Mnemiopsis* Shakers and found that they encode a functionally diverse array of voltage-gated K^+^ conductances with functional orthologs for many classic Shaker family subtypes found in cnidarians and bilaterians. Ctenophores therefore appear to have independently evolved much of the voltage-gated K^+^ channel diversity that is shared between cnidarians and bilaterians.

## Introduction

Voltage-gated K^+^ channels participate in a wide diversity of animal physiological systems but are most noted for shaping and patterning electrical responses in neurons and other excitable cells. Animal voltage-gated K^+^ channels display a high degree of functional diversity and are encoded by three structurally unique families: Shaker (Kv1-6, 8, 9) Ether-a-go-go (EAG, Kv10-12), and KCNQ (Kv7). All these gene families encode subunits that share a common core of six transmembrane domains (S1-S6), comprised of a voltage-sensor domain (VSD) and a K^+^-selective pore domain (PD) (Long et al., 2005; Whicher and MacKinnon, 2016; Sun and MacKinnon, 2017). Functional channels assemble as tetramers with a single pore surrounded by four independent VSDs. What structurally distinguishes the three gene families are unique family-specific intracellular domains that regulate gating and assembly. Shaker family channels have an N-terminal cytoplasmic T1 domain based on the BTB/POZ fold that self-tetramerizes and governs assembly (Pfaffinger and DeRubeis, 1995; Kreusch et al., 1998). In contrast, the cytoplasmic N-terminal EAG domain of EAG family channels has homology to Per-Arnt-Sim (PAS) domains (Morais Cabral et al., 1998) and works in tandem with the cytoplasmic C-linker/cyclic nucleotide binding homology domain (CNBHD) to regulate gating (Gustina and Trudeau, 2011; Gianulis et al., 2013; Whicher and MacKinnon, 2016; Wang and MacKinnon, 2017) and assembly (Lin et al., 2014). KCNQ family channels lack these domains but instead have a unique C-terminal coiled-coil domain that regulates channel assembly (Schwake et al., 2006; Sun and MacKinnon, 2017). The Shaker family encodes a broad diversity of depolarization-gated K^+^ currents including classic neuronal A-currents and delayed rectifiers. KCNQ family channels require PIP2 for gating and encode neuronal M-currents that regulate action potential threshold (Wang et al., 1998). The neurophysiology of the EAG family is less well characterized, but EAG family channels tend to activate with low voltage thresholds and influence subthreshold excitability (Saganich et al., 1999; Saganich et al., 2001; Zou et al., 2003; Zhang et al., 2009; Zhang et al., 2010; Kazmierczak et al., 2012; Li et al., 2015c; Hermanstyne et al., 2023).

Phylogenetic comparisons of diverse bilaterian and cnidarian species indicate that eight voltage-gated K^+^ channel types spanning these three gene families were present in the last common ancestor of Bilateria and Cnidaria (Jegla et al., 1995; Jegla and Salkoff, 1997; Jegla et al., 2009; Jegla et al., 2012; Martinson et al., 2014; Li et al., 2015b; Li et al., 2015c; Lara et al., 2023). The cnidarian/bilaterian Shaker family is comprised of four functionally independent gene subfamilies (Shaker or Kv1, Shab or Kv2, Shaw or Kv3 and Shal or Kv4) (Salkoff et al., 1992), while the cnidarian/bilaterian EAG family has three functionally independent gene subfamilies (Eag or Kv10, Erg or Kv11, and Elk or Kv12) (Warmke and Ganetzky, 1994). The KCNQ family has two sub-types widely conserved in bilaterians (Jegla et al., 2009), but only one ancestral lineage shared between cnidarians and bilaterians (Li et al., 2015b). Surprisingly, the voltage-gated K^+^ channel complement of the last common cnidarian ancestor appears to have been far more extensive than that of the last common bilaterian ancestor. The entire complement of bilaterian voltage-gated K^+^ channels is predicted to have descended from only eight to nine voltage-gated K^+^ channels present in the last common bilaterian ancestor (one from each EAG/Shaker subfamily and possibly two KCNQs) (Jegla et al., 2009; Li et al., 2015b; Li et al., 2015c; Lara et al., 2023). In contrast, extant cnidarians share 28 voltage-gated K^+^ channel gene lineages, with most of the diversification occurring within the Shaker family (Lara et al., 2023). Despite this large number of ancestral voltage-gated K^+^ channels, all cnidarian voltage-gated K^+^ channel families and subfamilies are shared with bilaterians.

Functional phenotypes of voltage-gated K^+^ channels are highly conserved between cnidarians and bilaterians (Jegla et al., 1995; Jegla and Salkoff, 1997; Jegla et al., 2012; Li et al., 2015b; Li et al., 2015c). However, cnidarian-specific gene duplications within the Kv1 and Kv4 subfamilies do expand the functional phenotypes of these subfamilies beyond what is typically seen in bilaterians. For example, bilaterian Kv4 subfamily channels all seem to encode classic A-type currents with closed state inactivation, whereas the large cnidarian Kv4 subfamily encodes functional orthologs of the bilaterian Kv4 channels, fast-inactivating Kv1-like currents and delayed rectifiers (Jegla and Salkoff, 1997; Li et al., 2015b). Similarly, the large cnidarian Kv1 subfamily includes channels with highly diverse kinetics and a broad range of activation thresholds (Jegla et al., 2012). Thus, phylogenetic lineage alone might not be sufficient to predict precise gating phenotypes of voltage-gated K^+^ channels over long evolutionary times or in the face of large, lineage-specific gene expansions.

The eight voltage-gated K^+^ channel lineages shared between cnidarians and bilaterians do not appear traceable to the origins of the nervous system. Recent genomic analyses place ctenophores, or comb jellies, as a sister lineage to all other animals (Ryan et al., 2013; Moroz et al., 2014; Whelan et al., 2015; Whelan et al., 2017; Schultz et al., 2023). They can therefore provide insights into voltage-gated K^+^ channel diversity in the last common ancestor of all animals at a time when the nervous system was first evolving. Examination of the first ctenophore genome (*Mnemiopsis leidyi*) revealed a large set of voltage-gated K^+^ channel genes, but these represented only two channel lineages shared with cnidarians and bilaterians: Kv1-like Shaker family genes (> 40 genes) and the EAG family (2 genes forming an outgroup to the cnidarian/bilaterian Kv10-12 subfamilies) (Li et al., 2015b). Recently, Kv2-4-like Shaker family channels were discovered in choanoflagellates, the protozoan sister lineage of animals (Jegla et al., 2024), but the Kv2-4 lineage is surprisingly not found in *Mnemiopsis*. These two lineages, Kv1-like and Kv2-4-like, can be differentiated based on the unique presence of a highly conserved Zn^2+^ binding site in the T1 interface of Kv2-4-like channels (Bixby et al., 1999; Jahng et al., 2002). No *Mnemiopsis* Shaker family channels have the Kv2-4-like Zn^2+^ binding site.

Comparison of choanoflagellates and *Mnemiopsis* with cnidarians and bilaterians therefore suggests the last common animal ancestor had as few as three voltage-gated K^+^ channel lineages (Kv1-like, Kv2-4-like, EAG), one of which was lost in the lineage leading to *Mnemiopsis* (Li et al., 2015b; Jegla et al., 2024). This raises the possibility that the complexity of electrical signaling, which requires functionally diverse voltage-gated K^+^ channels, was low at the time ctenophores diverged from the rest of animals. But does this mean electrical signaling in ctenophore nervous systems lacks some of the features present in nervous systems of cnidarians and bilaterians? Or just that signaling complexity simply evolved independently in ctenophores? The large expansion of Kv1-like Shakers in *Mnemiopsis* provides an opportunity to gain insight into how much independent functional diversification of voltage-gated ion channels has occurred in ctenophores. However, the functional significance of this expansion to ctenophores is unclear because its evolutionary origins are unknown and only five of the channels have been functionally expressed. MlShak1 and MlShak2 do not form functional channels as homomultimers although they can heteromultimerize with a cnidarian Shaker (Li et al., 2015b). MlShak3-5, in contrast, do function as homomultimers and but encode fast (MlShak4,5) or slow (MlShak3) inactivating K^+^ channels that are highly similar to cnidarian and bilaterian Kv1 currents (Jegla et al., 2012; Simonson et al., 2024). Fast inactivation in MlShak4 and MlShak5 even occurs by the same N-type ball and chain mechanism first described for *Drosophila* Shaker (Hoshi et al., 1990; Zagotta et al., 1990).

Here we examine the evolutionary origins of voltage-gated K^+^ channel diversity in *Mnemiopsis* Shaker family expansion by phylogenetically comparing voltage gated K^+^ channel sets from diverse ctenophores and functionally characterize a broader phylogenetic representation of *Mnemiopsis* Shakers. We find that most of the *Mnemiopsis* Shaker and EAG family expansion predates the radiation of three major ctenophore lineages: Beroe, Mnemiopsis and Hormiphora. In contrast, we found no evidence for the KCNQ family, the Kv2-4 Shaker subfamilies or the Kv10-12 EAG subfamilies in any ctenophore species. Functional expression of 16 *Mnemiopsis* Shaker channels representing conserved ctenophore Shaker lineages revealed an unexpectedly broad diversity of gating phenotypes. These results suggest that ctenophores have indeed independently evolved a functionally diverse set of voltage-gated K^+^ channels, primarily via expansion of the Kv1-like lineage of Shaker gene family.

## Materials & Methods

### Sequence Collection

We identified voltage-gated K^+^ channel sequences in *Beroe ovata* (Vargas et al., 2024) and *Hormiphora californensis* (Schultz et al., 2021) protein predictions with BLASTP searches using *Mnemiopsis leidyi* (Shaker and EAG families) (Li et al., 2015b; Li et al., 2015c) and *Nematostella vectensis* (KCNQ, HCN) (Li et al., 2015b) family members as queries. Protein family identity was confirmed with reciprocal BLASTP searches against *N. vectensis* proteins. Reciprocal best BLAST strategies have been very effective for comprehensively identifying voltage-gated K^+^ channel sets from *Mnemiopsis*, cnidarians, placozoans and choanoflagellates (Jegla et al., 2012; Li et al., 2015b; Li et al., 2015c; Jegla et al., 2018; Lara et al., 2023; Jegla et al., 2024). We used a similar pipeline to search the transcriptome of the ctenophore *Euplokamis dunlapae* (Whelan et al., 2017), except we started with a TBLASTN search and performed a reciprocal BLASTP using the translated results. Sequences were used for phylogenetic analysis if they were > 90% complete with respect to conserved domains.

### Alignment and Phylogenetic Analyses

We made alignments from the ctenophore EAG and Shaker family sequences in MEGA 11 (Tamura et al., 2021) using MUSCLE (Edgar, 2004) with default parameters. Gaps in conserved regions of a few gene predictions evident in these alignments were filled with sequence data from genomic and transcriptomic databases that were not included incorporated in the protein models. Transcriptomic and genomic data used to fill gaps 1) were confirmed to be on the same contig as the original predicted protein and 2) were located between the predicted sequences flanking the gaps. We removed poorly conserved linker regions with repeated independent length variations from the alignment and manually adjusted alignment around gaps for consistency prior to phylogenetic analysis. We included voltage-gated K^+^ genes from *Trichoplax adhaerens* (Placozoa), *Nematostella vectensis* (Cnidaria), *Drosophila melanogaster* (Bilateria, Protostomia), humans (Bilateria, Deuterostomia), and choanoflagellates (Shaker family only, *Salpingoeca dolithecata, Salpingoeca helianthica*, and *Mylnosiga fluctuans*) for comparison. We performed phylogenetic reconstruction of the channel families through Bayesian inference using MrBayes v3.2.7a (Ronquist et al., 2012) in combination with BEAGLE 3 (Ayres et al., 2019) under a mixed amino acid model. Phylogenies were generated with MCMC sampling for 3,000,000 generations using 2 independent runs of 6 chains sampled every 5,000 generations; the first 25% of samples were discarded to focus the analysis on the best trees. Standard deviations of split frequencies converged to 0.0003 and 0.001 for the EAG and Shaker phylogenies, respectively. We ran a second phylogenetic analysis for each alignment using maximum likelihood in IQ-TREE 2 (Minh et al., 2020) with an LG+F+R5 model (with 1000 ultrafast bootstrap replications) to corroborate results. To ascertain the number of voltage-gated K^+^ channels in the last common ancestor of *Beroe*, *Mnemiopsis*, and *Hormiphora*, we identified clades with strong statistical support (posterior probability > 0.95 for Bayesian phylogenies or bootstrap support > 0.95 for the maximum-likelihood phylogeny) that contained at least one sequence from *H. californensis* and at least one sequence from either *B. ovata* or *M. leidyi*. Statistically supported clades containing *E. dunlapae* sequence(s) along with at least one sequence from the other three species were inferred to have been present in the last common ancestor of these lineages. Because the *E. dunlapae* transcriptome almost certainly represents an incomplete subset of genes in the species, this study is unable to comprehensively predict all channel clades that are traceable to this early ctenophore ancestor.

### Functional Expression

Coding sequences for 14 *Mnemiopsis* Shaker genes (MlShak3-16) were synthesized using *Xenopus laevis* optimized codons (Twist Bioscience, CA, USA) and cloned into the pET21(+) vector (Twist Bioscience, CA, USA) between the EcoRI and NotI sites. We included 5’ and 3’ Xenopus β-globin UTR sequences from the pOX oocyte expression vector surrounding the ORFs to further optimize expression in oocytes (Jegla and Salkoff, 1997). The MlShak13 Δ2-18 transcription template was generated by PCR from the MlShak13 WT expression plasmid. Capped run-off transcripts were synthesized either from NotI-linearized templates using the mMESSAGE mMACHINE™ T7 Transcription Kit (ThermoFisher Scientific, Waltham, MA) or from PCR templates using 100U T7 polymerase (Takara, San Jose, CA), 4 mM CleanCap-AG (Trilink Biotechnologies, San Diego, CA), 5 mM each nucleotide, 5 mM DTT, 20U rnase inhibitor and 0.02U yeast inorganic pyrophosphatase (NEB, Ipswich, MA). RNA synthesis using the latter method, which we switched to part way through this study, was cheaper, more reliable and increased protein expression in oocytes (though relative expression differences between constructs were maintained); it is our recommended method.

*X. laevis* oocytes were defolliculated using 0.5-1 mg/mL type II collagenase (Sigma-Aldrich, St. Louis, MO) in calcium-free ND98 solution and cultured in ND98 with 2 mM Ca^2+^ supplemented with 100 U/mL/100 µg/mL/50 µg/mL of penicillin/streptomycin/tetracycline and 2.5 mM Na-Pyruvate (Li et al., 2015a; Simonson et al., 2024). Optimal RNA injection amounts were empirically determined and ranged from ∼ 0.05 to 25 ng/oocyte. Oocytes were incubated for 1-4 days at ∼18°C and then two-electrode voltage-clamp recordings were made under constant flow of low chloride solution (98mM Na^+^, 2mM K^+^, 2mM Cl^-^. 5mM HEPES, titrated to pH 7.2 with methanesulfonic acid) to reduce native Cl-currents (important when recording small currents (Baker et al., 2015)). Electrodes were made from borosilicate glass (Sutter Instrument, Novato, CA) and filled with 3mM KCl to tip resistances of ∼ 0.4-1.5 MΩ. Bath pellet electrodes were separated via a 1M NaCl 1% agarose bridge. Recordings were made using pClamp10/Digidata 1440a digitizer (Molecular Devices, Sunnyvale, CA), and a CA-1B amplifier (Dagan Instruments, Minneapolis, MN) run in TEV mode (ThermoFisher Scientific, Waltham, MA). Data were low pass filtered at 2-5 kHz and digitized at 4-10 kHz. We analyzed the data using OriginLab (Northampton, MA) and Clampfit (Molecular Devices). Boltzmann distributions were fit using the equation *f(V) = A_2_ + (A_1_ − A_2_)/[1/(1+e^(V−V_50_)/s^)]* where *s* is the slope, *A_1_* and *A_2_* are the lower and upper bounds, and *V_50_* is the half-maximal activation/inactivation voltage. The inactivation and recovery time course of MlShak13 was fit with a single exponential function (*I_t_ = I_to_ + Ae^-t/τ^*) where *I_t_* is the current at time *t*, *I_to_* is the initial current, *A* is the amplitude, and *τ* is the time constant.

## Results

### Sequence identification and phylogenetic analysis

To explore the evolutionary diversification of voltage-gated K^+^ channels within ctenophores, we identified voltage-gated K^+^ channel sequences from two newly available high-quality ctenophore genomes and transcriptomes from *Beroe ovata* (Vargas et al., 2024) and *Hormiphora californensis* (Schultz et al., 2023) using a comprehensive BLAST search strategy (see Methods). These species are believed to have shared a common ancestor with *Mnemiopsis leidyi* (Lobata) ∼170-260 million years ago (Whelan et al., 2017) and provide insights into the ancestral channel set of these three major ctenophore lineages. We also mined the transcriptome of *Euplokamis dunlapae*, which is a member of a lineage that branched off from the rest of ctenophores an estimated 350 million years ago (Whelan et al., 2017). While the *E. dunlapae* transcriptome is unlikely to represent the complete genome, it can nevertheless provide some insight into an earlier time point in the evolution of these channels within ctenophores. The phylogenetic relationship between ctenophores, parahoxozoans and choanoflagellates is shown in Fig. 1. Ctenophora and Parahoxozoa contain all animal lineages in which nervous systems are found, and choanoflagellates are the closest protozoan relatives of animals. Sponges, which do not have nervous systems, have been eliminated for visual simplicity but are believed to be the sister lineage of Parahoxozoa (Dunn et al., 2008; Ryan et al., 2013; Moroz et al., 2014; Li et al., 2021; Schultz et al., 2023).

**Figure 1.**
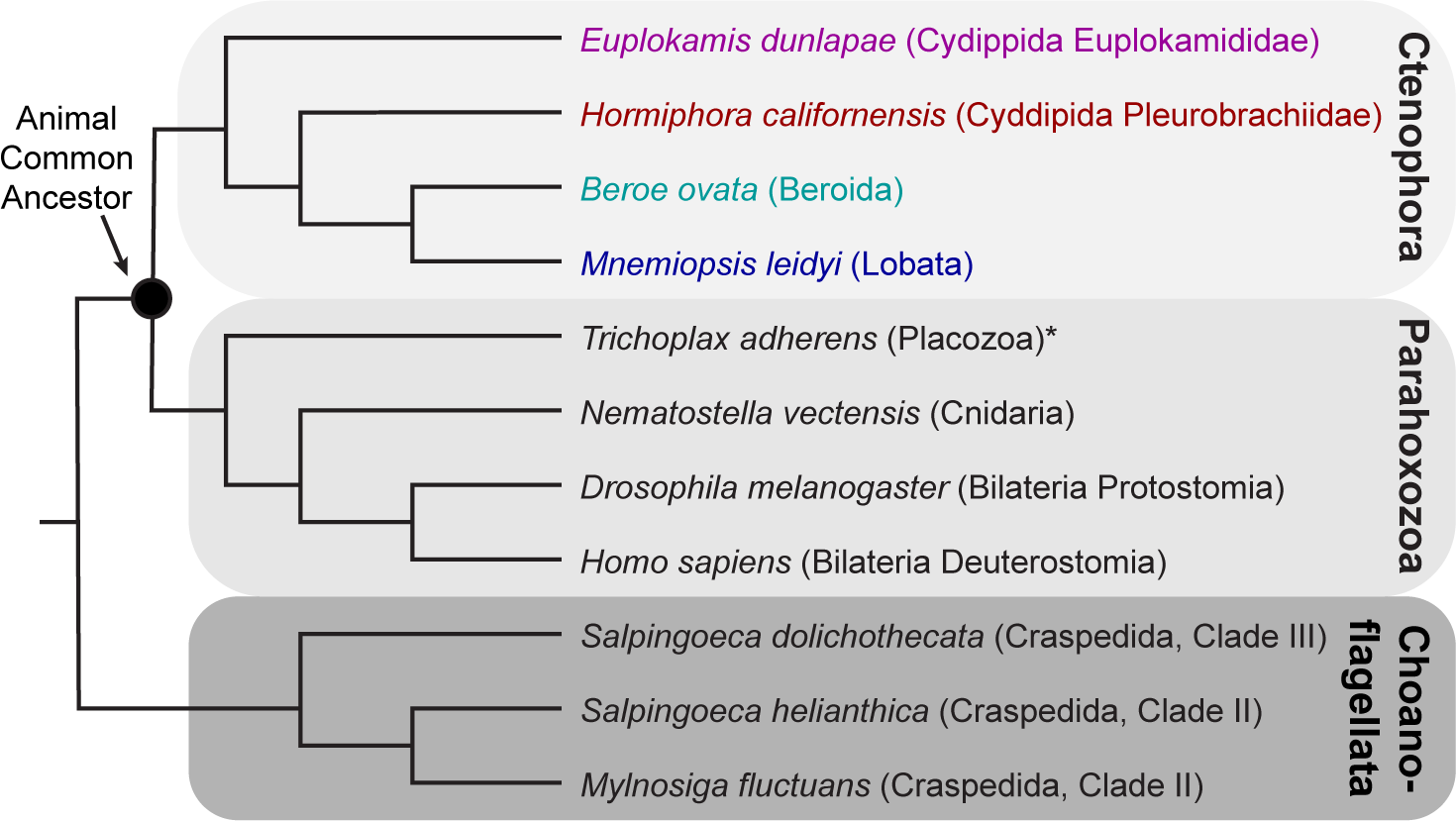
Phylogeny of animals and choanoflagellates. The schematic phylogeny depicts the relationships between animal and choanoflagellate species compared in our voltage-gated K^+^ phylogenies according to the most current view of the animal phylogeny. Ctenophora and Parahoxozoa are comprised of animals with nervous systems, except for the Placozoa (*). Sponges, which also do not have nervous systems, have been omitted for simplicity, but are now believed to be a sister group of Parahoxozoa based on chromosomal synteny (Schultz et al., 2023).

Voltage-gated K^+^ channel gene counts from four ctenophore species (*E. dunlapae, H. californensis, B. ovata and M. leidyi*) are compared to bilaterians (human, fly), a cnidarian (*Nematostella vectensis*), a placozoan (*Trichoplax adhaerens*) and three choanoflagellates (*Salpingoeca helianthica, Salpingoeca dolicothecata and Mylnosiga fluctuans*) in Table 1. Sequences for the *B. ovata*, *H. californensis* and *E. dunlapae* voltage-gated K^+^ channels described here are provided in File S1. These are lower bound estimates of the gene numbers, as we did not include a small number of incomplete Shaker family sequences, and the *E. dunlapae* transcriptome is unlikely to be comprehensive. We did not find any ctenophore KCNQ family genes and found no evidence for novel ctenophore-specific voltage-gated K^+^ channel families. The ctenophore EAG family was comprised of two ortholog pairs shared between *H. californensis* and *M. Leidyi* in our phylogenetic analyses (Table 1, Fig. 2), indicating that both genes were likely present in the last common ancestor of *Beroe*, *Mnemiopsis*, and *Hormiphora* despite their absence from the draft *B. ovata* genome. We used the HCN family of hyperpolarization-gated cation channels as an outgroup for the EAG family phylogeny since both families are members of the Cyclic Nucleotide Binding Domain (CNBD) superfamily of voltage-gated cation channels (Baker et al., 2015; Jegla et al., 2018). A single HCN channel has previously been reported for the cydippid ctenophore *Dryodora glandiformis* (Baker et al., 2015) and we show here that *Hormiphora californensis*, *Mnemiopsis leidyi* and *Beroe ovata*, also have one HCN channel (Table 1, Fig. 2, File S1). Ctenophore EAGs form an outgroup to the cnidarian/bilaterian Kv10-12 subfamilies as previously described (Li et al., 2015c). The EAG/HCN sequence alignment and tree file for Fig. 2 are provided as Files S2 and S3, respectively. We repeated the phylogenetic analysis using a maximum-likelihood approach (tree file provided as File S4) and reached the same conclusions: 1) ctenophore and parahoxozoan EAG family channels group separately and 2) there were two EAG channels in the last common ancestor of *Beroe*, *Mnemiopsis*, and *Hormiphora*. We did not find EAG or HCN family genes in *E. dunlapae*, but we can infer at least one of each must have been present in the ctenophore common ancestor from the presence of both gene families in ctenophores and parahoxozoans.

**Figure 2.**
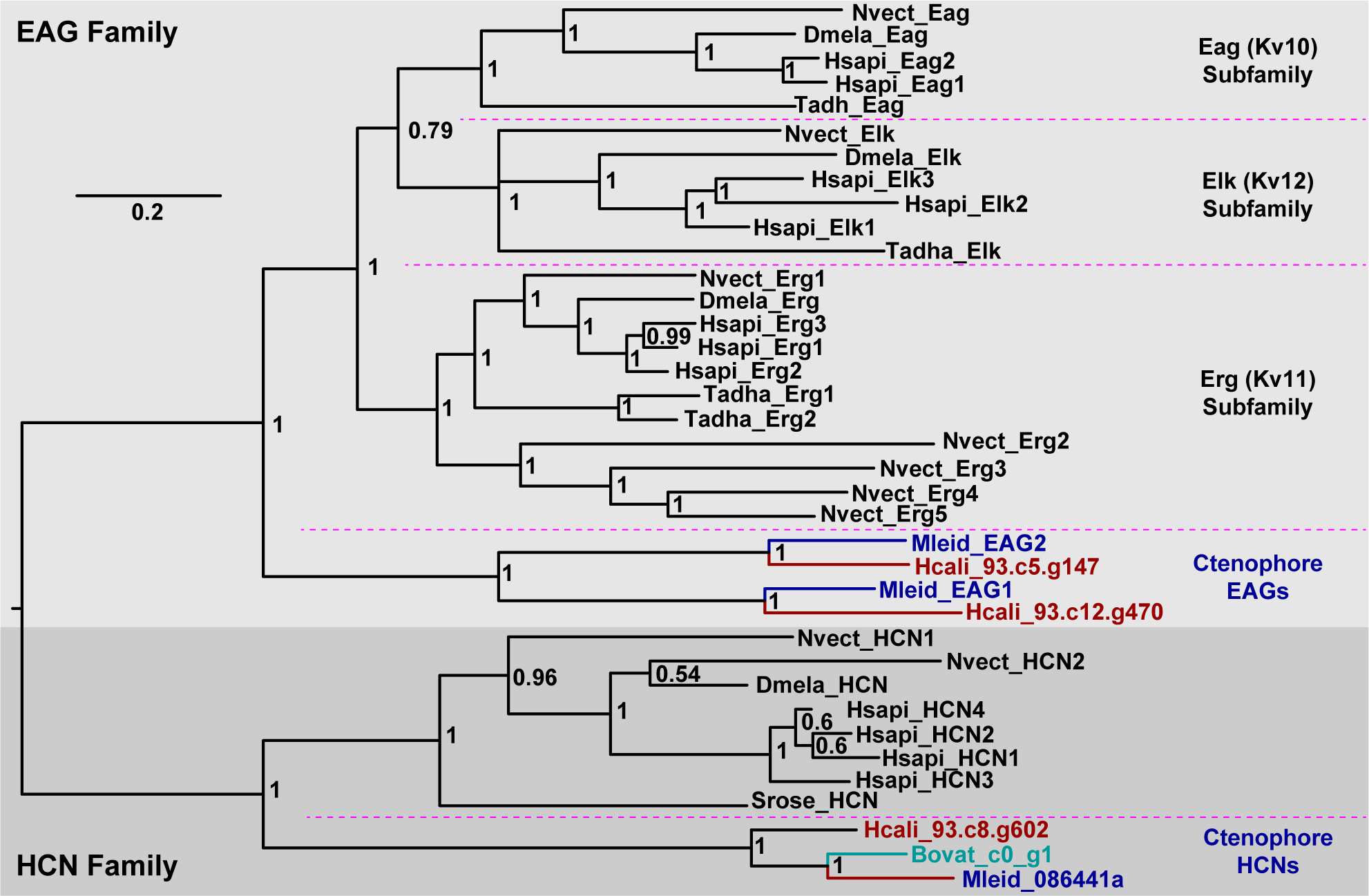
Bayesian inference phylogeny of the EAG and HCN voltage-gated ion channel channel families. Ctenophore sequences are color-coded according to the species phylogeny in Fig. 1, and gene names at branch tips use the following species prefixes: Hcali, *Hormiphora californensis*; Mleid, *Mnemiopsis leidyi*; Dmela, *Drosophila melanogaster*; Hsapi, *Homo sapiens*; Nvect, *Nematostella vectensis*; Srose, *Salpingoeca rosetta*; and Tadha, *Trichoplax adhaerens*. Numbers at nodes are posterior probability; > 0.95 indicate robust support. The scale bar for branch length is given in substitutions/site. The phylogeny was rooted between the EAG family (light gray shading) and HCN family (gray shading). Pink dotted lines are used to delineate the ctenophore clades and the parahoxoan Eag, Erg and Elk subfamilies within the EAG family. The sequence alignment used for the phylogeny and the tree file itself are provided as Files S2 and S3, respectively.

**Table 1.** Voltage-gated K^+^ channel and HCN channel gene numbers in select animals and choanoflagellate.

| Gene family | Shaker | EAG | KCNQ | HCN |
| --- | --- | --- | --- | --- |
| **Ctenophora** |  |  |  |  |
| *Beroe ovata* | 27 | 0 | 0 | 1 |
| *Euplokamis dunlapae* * | 11 | 0 | 0 | 0 |
| *Hormiphora californensis* | 33 | 2 | 0 | 1 |
| *Mnemiopsis leidyi* | 42 | 2 | 0 | 1 |
| **Placozoa** |  |  |  |  |
| *Trichoplax adhaerens* | 3 | 4 | 0 | 0 |
| **Cnidaria** |  |  |  |  |
| *Nematostella vectensis* | 45 | 7 | 1 | 2 |
| **Bilateria** |  |  |  |  |
| *Drosophila melanogaster* | 5 | 3 | 1 | 1 |
| *Homo sapiens* | 27 | 8 | 5 | 4 |
| **Choanoflagellata** |  |  |  |  |
| *Mylnosiga fluctuans* ** | 4 | 0 | 0 | 0 |
| *Salpingoeca dolichothecata* ** | 1 | 0 | 0 | 0 |
| *Salpingoeca helianthica* ** | 3 | 0 | 0 | 0 |
| *Salpingoeca rosetta* | 0 | 0 | 0 | 1 |

\*Inferred from transcriptome and therefore likely incomplete
\*\*Deep coverage transcriptome

In contrast to the EAG, KCNQ and HCN families, we identified numerous Shaker family channels in all ctenophore species, and the Shaker family comprises 112/116 of the voltage-gated K^+^ channels collectively found in *E. dunlapae, H. californensis, B. ovata and M. leidyi* (Table 1). Ctenophore Shaker gene numbers in the three genomes range from 27 in *B. ovata* up to 42 in *M. leidyi*, indicating expansion of the Shaker family is widespread ctenophores. In addition, we found 11 Shaker family channels in the draft transcriptome of *E. dunlapae* all of which form clades with at least one other ctenophore species. A Bayesian phylogeny of the Shaker gene family sequences from Table 1 is shown in Fig. 3 with the alignment and tree file provided as Files S5 and S6. The phylogeny indicates that much of the ctenophore Shaker family expansion predates the radiation of the *Beroe*, *Mnemiopsis*, and *Hormiphora*. We found 30 highly supported clades (posterior probability of 1) that contained sequences from *H. californensis* and at least one of *M. leidyi* and *B. ovata*, indicating that these 30 clades were present in the last common ancestor of *Beroe*, *Mnemiopsis*, and *Hormiphora*. We repeated the phylogenetic analysis using a maximum likelihood approach and recovered the same 30 ancestral clades (tree provided as File S6) with high (> 0.95) bootstrap support. In both phylogenies, the 11 *E. dunlapae* Shaker sequences fell into 9 different ancestral clades and one additional clade containing only *E. dunlapae* and *H. californensis* sequence. We interpret this clade as a 31^st^ ancestral channel clade that must have also been present in the last common ancestor of *Beroe*, *Mnemiopsis*, and *Hormiphora*. These *E. dunlapae* sequences indicate that much of the diversification of the Shaker family occurred early in ctenophore evolution, though the *E. dunlapae* data alone do not necessarily give us a comprehensive view of Shaker family and EAG family voltage-gated K^+^ channel genes present in the earliest ctenophores.

**Figure. 3.**
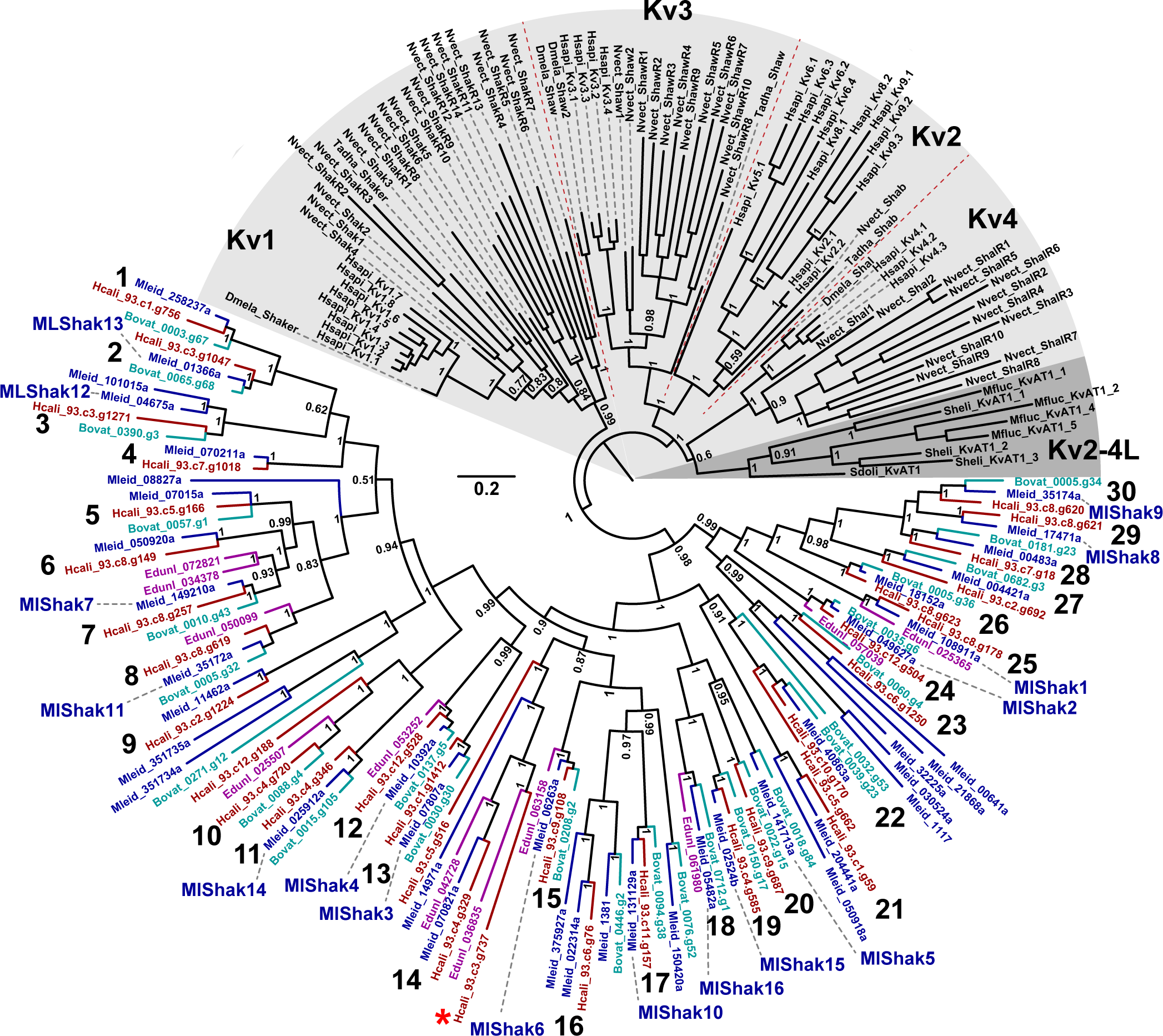
Bayesian inference phylogeny of the animal and choanoflagellate Shaker family of voltage-gated K^+^ channels. Ctenophore sequences are color coded as in previous figures and numbers at nodes indicate posterior probability and species prefixes follow the convention defined in Fig. 2 with the addition of three choanoflagellates: Mfluc, *Mylnosiga fluctuans*; Sdoli, *Salpingoeca dolichothecata*; and Sheli, *Salpingoeca helianthica*. The phylogeny is rooted for display between clades that do (Kv2-4, choanoflagellates) or do not (Kv1, ctenophore Shakers) contain the diagnostic T1 Zn^2+^ binding site, and a scale bar (substitutions/site) is given for branch length. Parahoxozoan (Kv1-4) and choanoflagellate (Kv2-4L) Shaker clades are shaded light gray and dark gray, respectively, and the four recognized parahoxozoan Shaker subfamilies are delineated with dotted red lines. Ancestral clades present in the last common ancestor of *Beroe*, *Mnemiopsis*, and *Hormiphora* are labelled 1-30, and 16 channels we expressed here are labelled MlShak1-16. A 31^st^ putative ancestral clade found in *H. californensis* and *E. dunlapae* is indicated with an asterisk. The alignment and tree are Files S5,6.

The ctenophore Shaker family sequences formed a single highly supported clade in unrooted phylogenies separate from all parahoxozoan and choanoflagellate sequences, indicating that the gene expansions observed in ctenophore species are fully ctenophore specific. The Shaker family phylogenies presented in Fig. 3 and File S6 contain three major clades: 1) the ctenophore clade, 2) the parahoxozoan Kv1 subfamily, and 3) a clade comprised of the parahoxozoans Kv2-4 subfamilies together with the choanoflagellate Kv2-4-like Shakers. Channels in the latter clade share the T1 Zn^2+^ binding site that clearly indicates a single common origin. This site is absent from all Kv1 subfamily channels and all ctenophore Shaker family channels, but the phylogeny does not conclusively resolve the relationship between ctenophore Shakers and the Kv1 subfamily (see Discussion).

### Functional expression

We did not attempt to functionally express ctenophore EAG or HCN channels in this study because they were not amenable to gene synthesis. We instead focused here on functional characterization of the Shaker family expansion that represents the bulk of the ctenophore voltage-gated K^+^ channel set. MlShak1-5, the previously expressed ctenophore Shakers, represent only a small fraction of the phylogenetic diversity of ctenophore Shakers shown in Fig. 3, and are thus unlikely to represent the broader functional diversity of ctenophore Shakers. We therefore synthesized codon-optimized *Xenopus* oocyte expression vectors for 11 additional *M. leidyi* Shaker genes from across the breadth of the phylogeny which we name MlShak6-16 here. These 11 genes were selected from the full gene set because 1) we had high confidence in the full coding sequence predictions, including sequence at the N- and C-termini, from transcriptome data and/or cross-species conservation, 2) they had coding sequence lengths compatible with efficient gene synthesis, and 3) they each represented one of the conserved Shaker channel clades present in the last common ancestor of *Beroe*, *Mnemiopsis*, and *Hormiphora*. MlShak1-16 collectively represent 15 of these ancestral clades (Fig. 3). Only the MlShak5 clade has a topology suggesting it may have arose after *Mnemiopsis* and *Beroe* split from *Hormiphora*, although it may only represent a loss of this family in *Hormiphora*. Sequences for the MlShak6-16 expression vectors are given in File S7.

Fig. 4a shows example traces recorded in response to step depolarizations from oocytes expressing MlShak1-16. MlShak1-5 current phenotypes shown here are consistent with previous reports (Li et al., 2015b; Simonson et al., 2024). A total of 9/16 channels (MlShak3-9, MlShak13 and MlShak15) consistently produced voltage-gated K^+^ currents (Fig. 4a,b) when expressed in isolation, and thus are able to form functional homomeric channels. We first injected RNAs at high concentration (∼10-25ng/oocyte) which in our historical lab experience with > 200 different ion channels, is sufficient to see even low efficiency functional channel formation in oocytes. We therefore interpret the absence of detectable K^+^ currents for MlShak1,2,10-12,14,16 (Fig. 4 a,b) to mean they are either unable to assemble as homomultimers or unable to gate by voltage alone. Each of these non-expressing channels was injected in multiple oocyte batches to confirm the phenotype, but Fig. 4b reports current size data from 1-2 batches. These channels may be equivalent to cnidarian/bilaterian silent or regulatory subunits that require heteromeric assembly for functional expression (Jegla and Salkoff, 1997; Jegla et al., 2012; Bocksteins, 2016; Pisupati et al., 2018; Jegla et al., 2024), but we did not test heteromeric combinations in this study. MlShak3-6 currents were among the best expressing channels we have observed in the oocyte system and the data reported in Fig. 4b was recorded from oocytes injected with ∼0.05-0.1 ng RNA/oocyte. What is immediately apparent from Fig. 4a is the diversity of gating phenotypes, ranging from rapid inactivation (MlShak4, MlShak5 and MlShak13) through slow inactivation (MlShak3 and MlShak15) to almost no inactivation during 400 millisecond (ms) test steps (MlShak6-9). Most of the channels displayed fast activation typical of bilaterian and cnidarian Shaker family channels, with exception of MlShak8 and MlShak9. These two channels required multiple seconds for full activation, a phenotype that has not previously been observed in the Shaker family to our knowledge. For MlShak9, activation was so slow it limited our ability to collect data at high voltages. We do not quantify kinetics for these channels in detail here (except for MlShak13 inactivation, below), but the properties of currents were extremely consistent from oocyte to oocyte.

**Figure 4.**
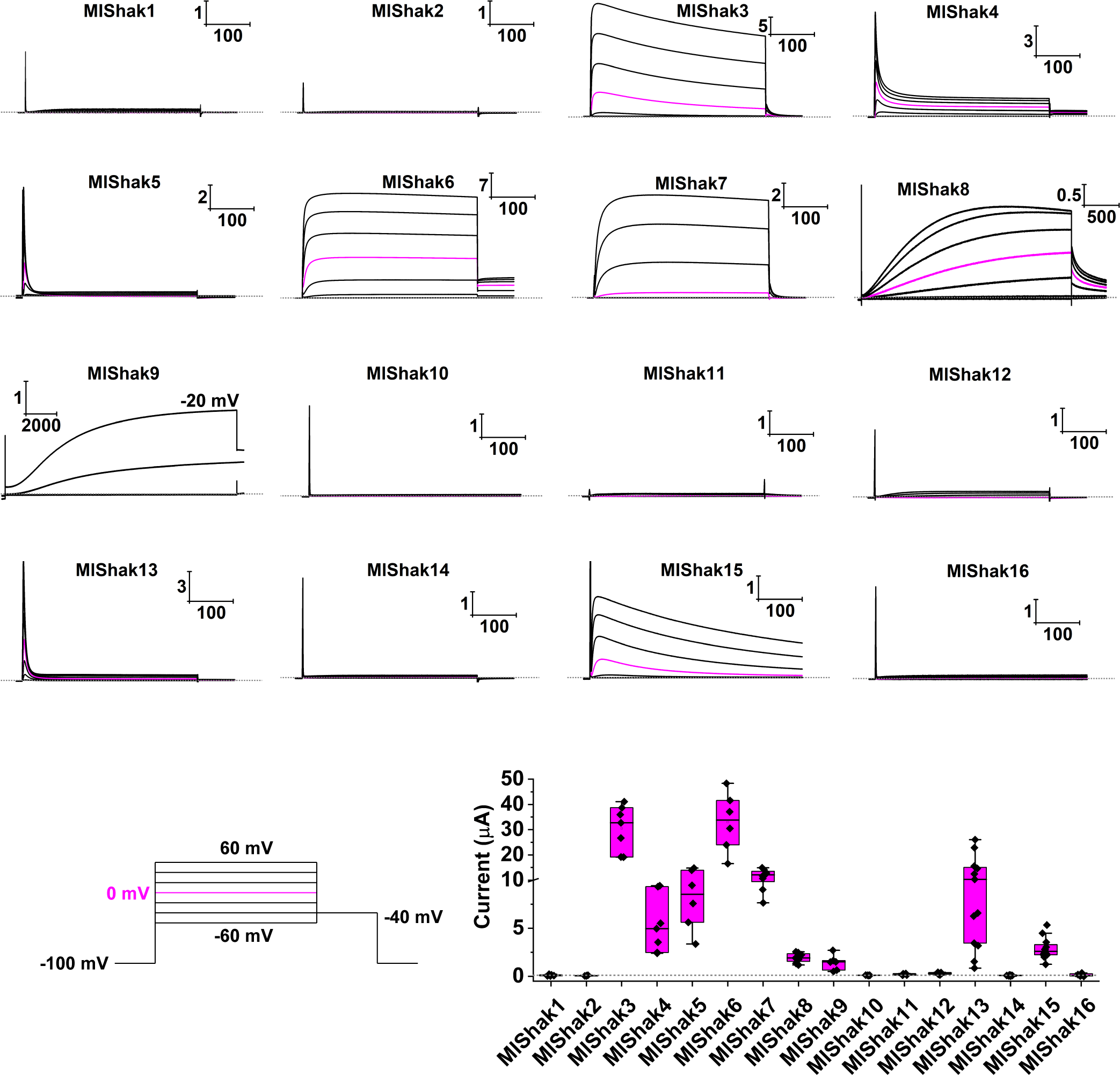
Functional expression of MlShak1-16 in *Xenopus* oocytes. (A) Example current traces families recorded in response to step depolarizations for MlShak1-16. Scale bars indicate current (μA) and time (ms), the 0 current level is indicated with a dotted line and the 0 mV traces is highlighted in pink to help visualize difference in activation thresholds. The inset at the bottom left indicates the standard voltage protocol, with 400 ms steps delivered every 20 mV from -60 to 60 mV. A -40 mV tail voltage was standard in these examples, but - 60 mV was used for MlShak6 and MlShak13, and -20 mV for MlShak7. MlShak5 and MlShak13 were held at -120 mV for recovery from inactivation and removal of steady state inactivation, respectively. MlShak10, 15 and 16 were recording using 1 s steps, so the end of the pulse is not visible in the 500 ms time window displayed. Note MlShak8 and MlShak9 activate very slowly and are displayed on much longer time scales. The top MlShak9 trace displayed here is -20 mV, so there is no pink 0 mV trace highlighted. 1 ms bracketing the capacitive transient peak was clipped from the traces for display purposes, but there are still small residual capacitive current spikes in some traces; these were left because they help mark the beginning and end of steps for channels that did not express. (B) Peak current size at +60 mV for representative oocyte batches expressing MlShak1-16. Note the Y-axis scale changes at the slashed line to allow visualization of high and low expressing channels on the same plot. Box plot shading indicates the 25^th^ and 75^th^ percentile, whiskers mark the data range, the line indicates the median, and black diamonds are individual data points. N numbers were in order: 7, 5, 7, 7, 6, 6, 8, 8, 6, 5, 4, 6, 13, 8, 11, 12, 7. No exogenous voltage-gated K^+^ currents were detected for MlShaks 1,2, 10, 11, 12, 14 and 16.

Rapid inactivation in MlShak4 and MlShak5 occurs by a classic N-type ball and chain mechanism shared with *Drosophila* Shaker (Simonson et al., 2024). Here we identify MlShak13 as a third rapidly inactivating ctenophore Shaker and tested it for N-type inactivation by expressing a truncated version lacking the native N-terminus (MlShak13 Δ2-18). This truncation removed fast inactivation from the channel implicating the same classic N-type inactivation mechanism (Fig. 5a). The MlShak13 inactivation time course is similar to rapid inactivation in MlShak4 and MlShak5 in that all have time constants faster than 6 ms above 0 mV (Fig. 5b and (Simonson et al., 2024)). However, MlShak13 differs markedly from MlShak4 and MlShak5 in having rapid recovery from inactivation (Fig. 5c) with a time constant of 35.5 ± 0.7 ms at -120 mV (n = 4) compared to 552 ± 116 ms and 2822 ± 452 ms previously reported for MlShak4 and MlShak5 (Simonson et al., 2024), respectively.

**Figure 5.**
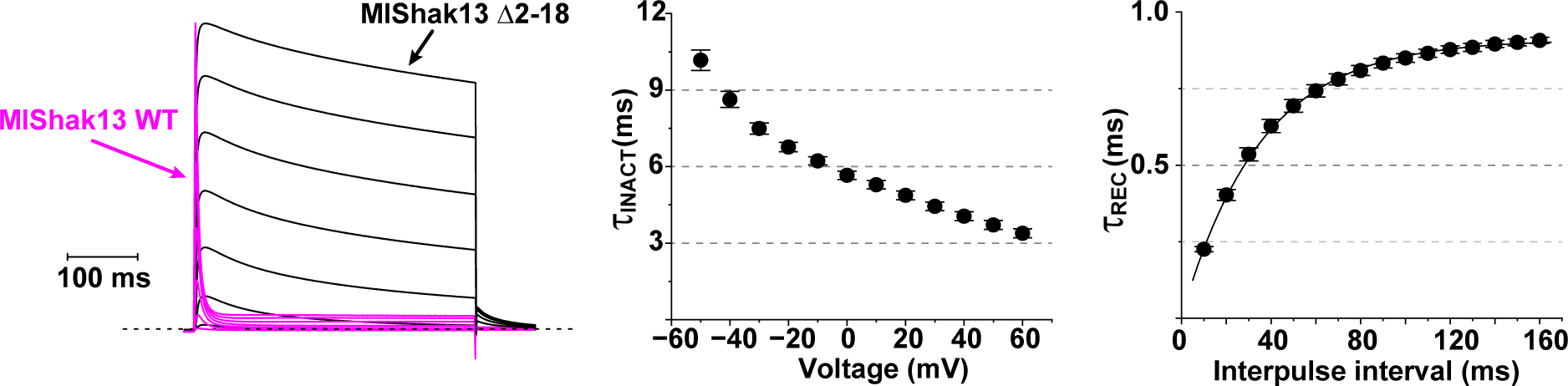
MlShak13 uses an N-type inactivation mechanism. (A) Families of outward K^+^ currents recorded from an oocyte expressing MlShak13 WT (pink, replotted from Fig. 4A) and MlShak13 Δ2-18 (black) show that the N-terminus confers fast inactivation. Currents were normalized to the peak value at +60 mV for comparison, capacitive transients have been clipped from the traces as described in Fig. 4, and a scale bar is given for time. Currents were recorded in response to 400 ms depolarizations ranging from -60 to +60 mV in 10 mV increments and tails were measured at -60 mV. (B) Time constant of fast inactivation for MlShak13 WT for the indicated voltages for single exponential fits of the inactivation time course. Data show mean ± S.E.M., N = 10. (C) Time constant of recovery from inactivation determined using a double pulse protocol in which two 400 ms pulses to 40 mV were separated by a recovery step to -120 mV of the indicated time. Data show mean ± S.E.M. (N = 4) peak current recorded in the 2^nd^ test step normalized to the peak current of the first test step. The smooth curve is a single exponential fit with a time constant of 35.5 ms and peak recovery of 0.91.

We generated conductance-voltage curves (GV) from isochronal tail currents for the 9 MlShak channels that expressed as homomultimers to compare their voltage-activation ranges (Fig. 6a). Average V_50_ and slope values calculated from single Boltzmann fits of data from individual eggs are shown in Table 2 and were used to generate the smooth curves in Fig. 6a. MlShak3-5 GV data in Table 2 were previously reported (Simonson et al., 2024) and the curves shown in Fig. 6a for comparison were generated with the V_50_ and slope values from that study. Data for all three fast inactivating channels (MlShak4,5 13) were generated using N-terminal truncated constructs that remove fast N-type inactivation to reveal tail currents. The V_50_ values of the GVs for the 9 MlShak homomultimers spanned a range of ∼ 65 mV from -45.7 ± 0.8 mV in MlShak9 to 18.9 ± 1.1 mV in MlShak7. This range is comparable to the ∼58 mV V_50_ range for GVs of all reported homomeric cnidarian Kv1-4 channels combined (Jegla et al., 1995; Jegla and Salkoff, 1997; Bouchard et al., 2006; Sand et al., 2011; Jegla et al., 2012; Li et al., 2015b). We also generated steady state inactivation (SSI) curves with 6 second (s) preconditioning steps prior to a depolarizing test step used to measure available current. SSI data are shown with single Boltzmann fits in Fig. 6b. SSI fit parameters are reported in Table 2 and V_50_ values ranged from -105.8 ± 0.7 mV in MlShak13 to -28.5 ± 1.1 mV in MlShak7. This range of SSI V_50_s (∼77 mV) is again comparable to the entire range of SSI V_50_s reported for cnidarian Shaker family channels spanning the Kv1-4 subfamily (∼73 mV) (Jegla et al., 1995; Jegla and Salkoff, 1997; Bouchard et al., 2006; Jegla et al., 2012; Li et al., 2015b).

**Figure 6.**
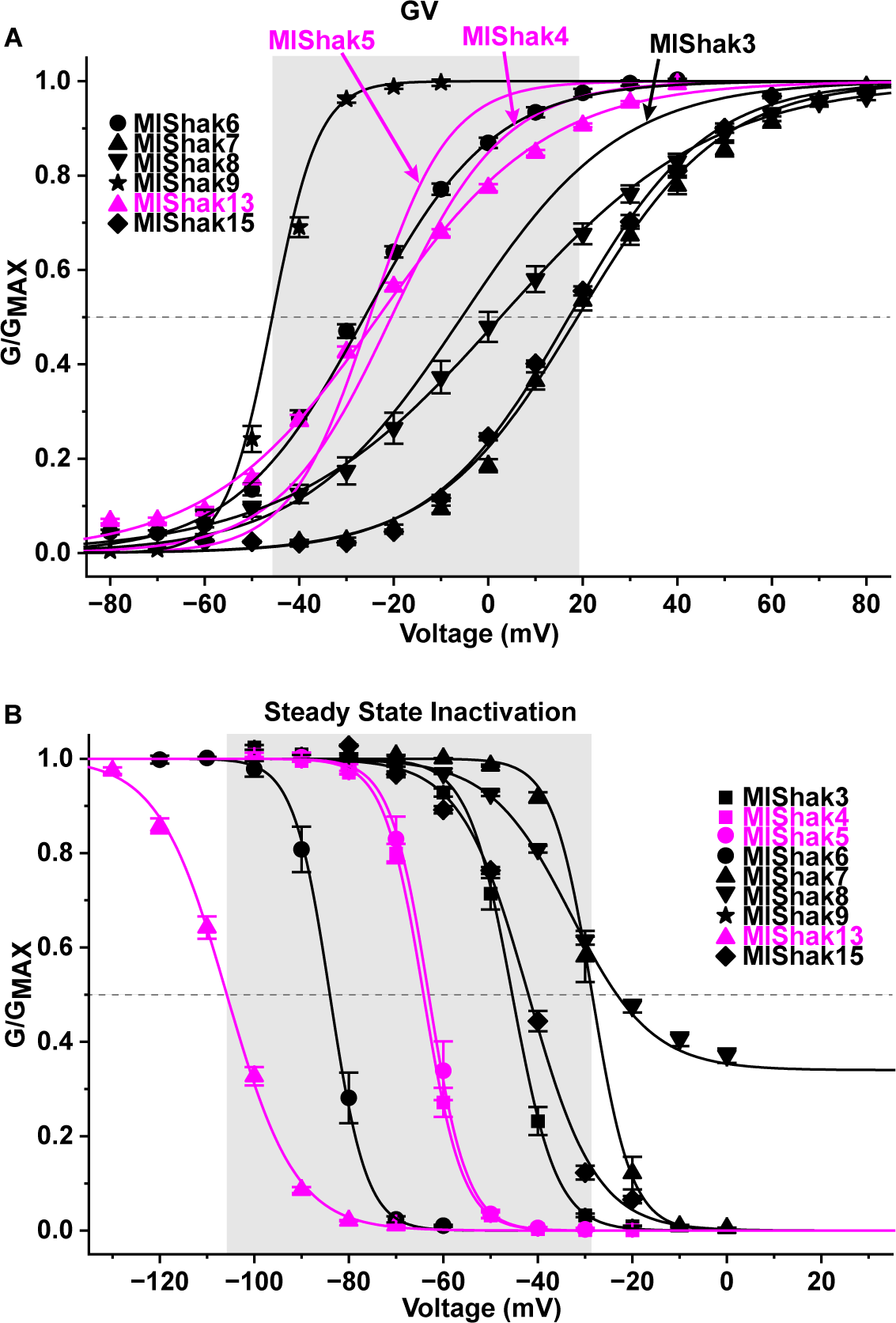
(A) Conductance voltage (GV) and (B) Steady state inactivation (SSI) curves for homomeric MlShak channels. Channels with fast N-type inactivation are highlighted in pink, and truncated constructs lacking fast inactivation were used for GV determination. Data points show mean ± S.E.M. (N given in Table 2), and data are normalized for comparison. Smooth curves show single Boltzmann fits of the data using the mean V_50_ and slope values from fits of individual oocytes (Table 2). Only GV curves are shown for MlShak3-5 because their data has been previously reported (Simonson et al., 2024), and the curves were made using V_50_ and slope values from that study. The gray dotted line marks G/Gmax = 0.5 and the gray shading marks the voltage range between the highest and lowest V_50_ values. GV data was taken from isochronal tail currents recorded at -60 mV (MlShak6,13), -40 mV (MlShak8,9,15), -20 mV (MlShak3-5) or 0 mV (MlShak7). Test pulse lengths were 50 ms (MlShak13 Δ2-18, MlShak15), 100 ms (MlShak7), 400-500 ms (MlShak3-6), 3 s (MlShak8) or 15 s (MlShak9), and were determined based on activation time course and inactivation properties. The holding potential was -100 mV for all except MlShak13 Δ2-18 (-120 mV). SSI data was recorded by measuring isochronal currents in test steps to +40 mV after 5-6 s pre-conditioning steps to the indicated voltages. Holding potentials were -100 mV except for MlShak5 and MlShak13 (-120 mV).

**Table 2.** V_50_ and Slope values for single Boltzmann fits of MlShak GV and SSI data.

|  | GV fits |  |  | SSI fits |  |  |
| --- | --- | --- | --- | --- | --- | --- |
| | $V_{50}$ (mV) | Slope (mV) | N | $V_{50}$ (mV) | Slope (mV) | N |
| MIShak3* | $-5.6 \pm 1.3$ | $16.8 \pm 0.35$ | 10 | $-45.4 \pm 0.8$ | $4.66 \pm 0.13$ | 8 |
| MIShak4* | $-20.5 \pm 0.85$ | $11.9 \pm 0.27$ | 11 | $-64.2 \pm 0.6$ | $4.16 \pm 0.07$ | 6 |
| MIShak5* | $-25.1 \pm 0.33$ | $8.4 \pm 0.27$ | 9 | $-63.1 \pm 1.3$ | $4.06 \pm 0.20$ | 4 |
| MIShak6 | $-26.2 \pm 1.4$ | $13.7 \pm 0.6$ | 6 | $-84.0 \pm 1.2$ | $2.86 \pm 0.14$ | 6 |
| MIShak7 | $18.9 \pm 1.1$ | $15.2 \pm 0.5$ | 8 | $-28.5 \pm 1.1$ | $4.14 \pm 0.13$ | 7 |
| MIShak8 | $2.6 \pm 2.7$ | $22.4 \pm 1.2$ | 7 | $-33.0 \pm 0.4$ | $8.55 \pm 0.18$ | 6 |
| MIShak9 | $-45.7 \pm 0.8$ | $4.89 \pm 0.2$ | 6 | N.D. | N.D. | |
| MIShak13 | $-21.1 \pm 0.6$ | $14.5 \pm 0.4$ | 7 | $-105.8 \pm 0.7$ | $7.29 \pm 0.17$ | 6 |
| MIShak15 | $17.1 \pm 0.6$ | $14.5 \pm 0.43$ | 8 | $-42.1 \pm 0.44$ | $7.15 \pm 0.34$ | 6 |

Data are given as Mean ± S.E.M. of N oocytes.
\*GV data taken from (Simonson et al., 2024)
N.D. (not done)

## Discussion

### Evolution of animal voltage-gated K^+^ channels

Here we show that the large set of voltage-gated K^+^ channel genes are a common feature of ctenophores, the earliest diverging animal lineage. Figure 7 shows a comparison of our current view of the ancestral voltage-gated K^+^ channel sets for bilaterians, cnidarians and ctenophores, the animals with nervous systems. It is remarkable that ctenophores and cnidarians, whose nervous systems are often described as simple relative to bilaterian animals, have much larger predicted ancestral channel sets than bilaterians. This suggests that a large voltage-gated K^+^ channel set was probably not a requirement of the origin of centralized nervous systems. It may be sufficient that the ancestral bilaterian channel set represents a functionally diverse group of independent (non-mixing) channel types. The ctenophore K^+^ channel set, while large, is markedly different than the K^+^ channel sets found in cnidarians and bilaterians in that it derives from just two ancestral lineages rather than eight (Jegla et al., 2009; Li et al., 2015b; Lara et al., 2023). Our analyses suggest that early ctenophores (at least as old as the last common ancestor of Mnemiopsis and Hormiphora) had a complement of at least 31 Shaker family channels and 2 EAG family channels (Fig. 7). For both of these families, the ctenophore channels are more closely related to each other than they are to any of the cnidarian and bilaterian channels. This suggests that single EAG and Shaker family channels in the last common ancestor of all animals independently radiated in each of these lineages to give rise to the full diversity of these channels in modern ctenophore and non-ctenophore animals. This same evolutionary pattern is observed in innexin channels (Ortiz et al., 2023). The cnidarian/bilaterian KCNQ family is entirely absent in ctenophores and the ctenophore Shaker and EAG families lack the 7 functionally independent cnidarian/bilaterian subfamilies. KCNQs and these Shaker/EAG subfamilies might have evolved after the divergence of ctenophores and parahoxozoans. All are also absent to date from sponges and KCNQ channels and the Shaker Kv4 subfamily are missing in placozoans (Li et al., 2015b).

**Figure 7.**
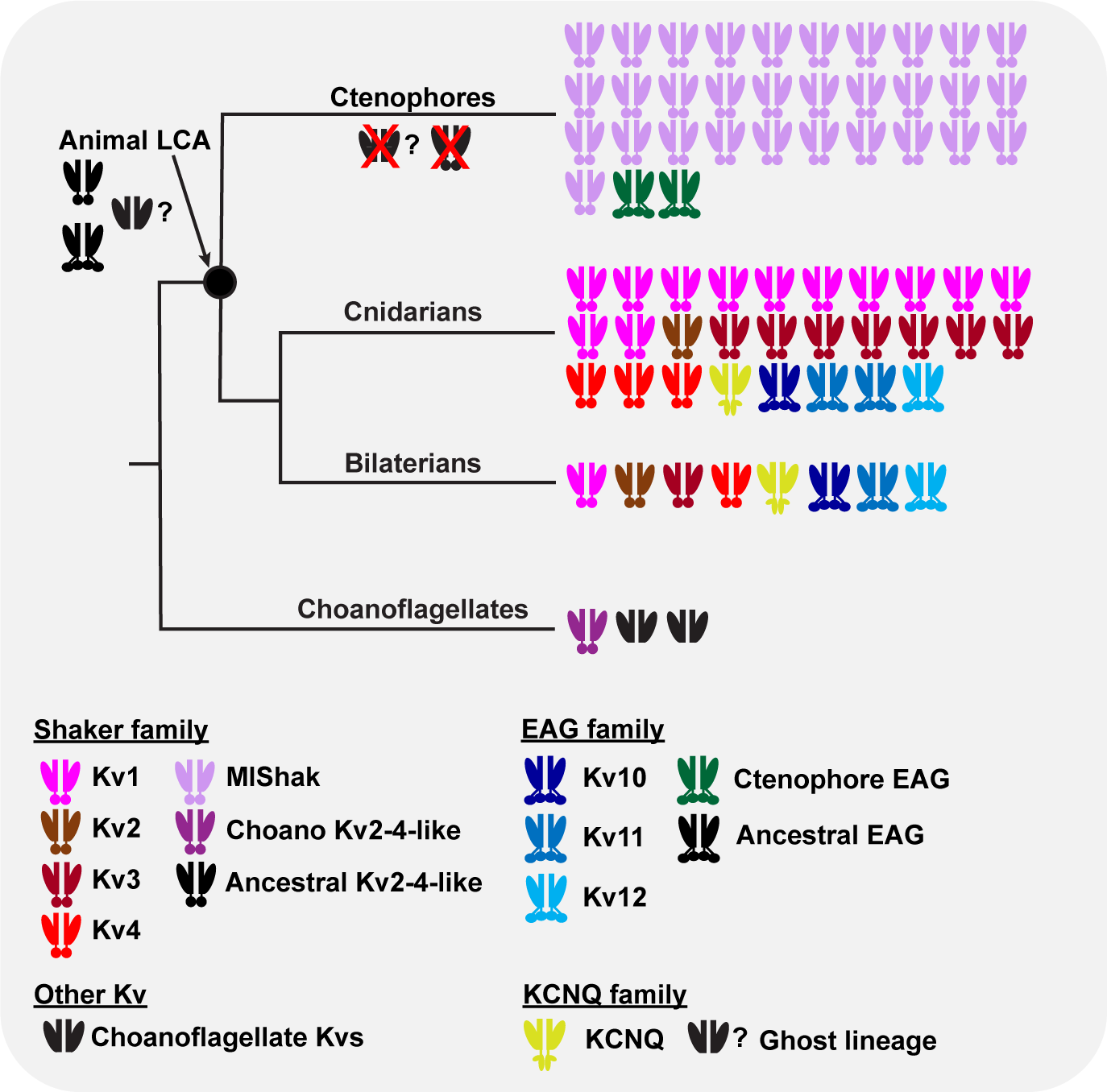
Ancestral voltage-gated K^+^ channel (Kv) sets for bilaterians, cnidarians, ctenophores and choanoflagellates. Only animals with nervous systems are depicted. Each icon represents a single ancestral gene inferred from phylogenetic comparison of channels from multiple living species. Icons are shaped colored by gene family and subfamily (or clade) according to the legend at the bottom. The ctenophore channel set defined here corresponds to the ancestor of Mnemiopsis, Beroe and Hormiphora. Almost the entire ctenophore Kv set has evolved from a single Kv1-like Shaker family lineage, whereas cnidarians and bilaterians share eight independent lineages spanning three gene families. Ctenophores appear to have independently evolved a high degree of biophysical diversity within their single Shaker lineage. The Kv set inferred for the animal last common ancestor (LCA) from comparison of animals and choanoflagellates is also depicted and a question mark indicates uncertainty about the presence of a KCNQ ghost lineage. This lineage, if present, and the ancestral Kv2-4-like Shaker lineage, have been lost in ctenophores (red Xs). Here we assume independent origins for Ctenophore Shakers and the Kv1 subfamily based on phylogeny (Fig. 3), but if they instead stem from a single T1 Zn^2+^ binding site loss event, then a fourth Kv1-like lineage could have been present in the animal LCA.

Given that EAG channels have not been identified in choanoflagellates or other non-animals (Jegla et al., 2024), it is likely that the this family arose in the animal stem after the split of choanoflagellates and animals. However, the absence of EAG channels in protozoans should be revisited as more species are sequenced to rule out selective loss in choanoflagellates. Cyclic Nucleotide Binding Domain (CNBD) superfamily cation channels (to which the EAG family belongs) are widespread across the tree of life (Schachtman et al., 1992; Sentenac et al., 1992; Jegla and Salkoff, 1994, 1995; Brams et al., 2014; Jegla et al., 2018; Pozdnyakov et al., 2020) but the relationships between these channels in the major eukaryotic lineages have not yet been defined (Jegla et al., 2018; Pozdnyakov et al., 2020). The ancestral lineage from which the KCNQ family evolved is also not clear because there are no orphan voltage-gated K^+^ channels in early animals that fall outside the EAG and Shaker lineages. It is possible KCNQs evolved from Shakers via loss of T1 because both share a domain-swapped VSD arrangement not found in EAGs (Long et al., 2005; Whicher and MacKinnon, 2016; Sun and MacKinnon, 2017), but there isn’t a smoking gun of sequence homology to favor this origin. Alternatively, they might have evolved from a ghost lineage of voltage-gated K^+^ channels that was lost in the early diverging animal lineages. This would be analogous to the Shaker Kv2-4-like lineage that we can infer was present in the animal last common ancestor only from its presence in Choanoflagellates (Jegla et al., 2024), the protozoan sister clade to animals. Choanoflagellates do have two voltage-gated K^+^ channel lineages in addition to Shaker (Jegla et al., 2024), but these have not been phylogenetically compared to KCNQs and there is as yet no evidence for their presence in animals.

We call the large ctenophore Shaker family channels here Kv1-like because, like the bilaterian/cnidarian Kv1 subfamily, they lack the T1 Zn^2+^-binding site shared between the animal Kv2-4 and choanoflagellate Shakers (Bixby et al., 1999; Jegla et al., 2024). Given our phylogeny, it is likely that these two subfamilies each independently lost the T1 Zn^2+^-binding site and we therefore cannot be certain of the presence of Kv1-like channels in the animal last common ancestor. Ctenophore Kv1-like channels span a broad range of biophysical properties comparable to the Shaker families of cnidarians and bilaterians, including functional orthologs of classic delayed rectifiers (MlShak6), rapid N-type inactivating Shakers (MlShak4, 5, 13), and suites of channels tuned for activity at hyperpolarized (MlShak4-6, 9, 13) and depolarized (MlShak7,15) voltages, as found in both cnidarians and vertebrates (Jegla et al., 1995; Jegla and Salkoff, 1997; Jegla et al., 2012; Li et al., 2015b). We examined only half of the conserved ctenophore Shaker family lineages, so it is quite possible that the ctenophore Shaker family is even more functionally diverse than we show here. Because ctenophores and parahoxozoans share a broad array of biophysical phenotypes across their Shaker families, but share only one ancestral Shaker gene lineage, many of these biophysical similarities are likely to have evolved independently.

### Open questions regarding the functional diversity of ctenophore Shakers

In both cnidarians and vertebrates, silent/regulatory subunits that require heteromeric assembly further diversify the functional properties of Shaker family channels. For example, the vertebrate silent/regulatory subunit Kv6.4 introduces closed state inactivation into Kv2 subfamily delayed rectifiers (Ottschytsch et al., 2002; Pisupati et al., 2018), and Kv4 subfamily silent/regulatory subunits in cnidarians diversify the voltage-activation range and inactivation kinetics of classic A-type currents (Jegla and Salkoff, 1997; Li et al., 2015b). In fact, more than half of all cnidarian Shaker family genes are likely to encode silent/regulatory subunits (Lara et al., 2023). Seven out of the 16 *Mnemiopsis* channels we expressed failed to form functional homomeric channels and are therefore potential silent/regulatory subunits. In support of this, MlShak1 and MlShak2 do form functional heteromultimers when co-expressed with a cnidarian Kv1 subfamily channel (Li et al., 2015b). However, proving the silent/regulatory phenotype requires finding native assembly partners in ctenophores, a potentially extensive combinatorial expression process that we did not undertake here. Alternatives for the lack of current include gating requirements other than voltage changes (rare for Shakers) or errors in the coding ORF predictions that block homomeric expression. The latter possibility is unlikely because our ORF predictions were supported by cross-species conservation. It is highly unlikely that these non-expressers are remnant pseudogenes because they represent conserved ancestral lineages.

One of the key underpinnings of complex electrical signaling in cnidarians and bilaterians is that their eight conserved voltage-gated K^+^ channel families and subfamilies are functionally independent because they are not assembly compatible. This allows for expression of up to eight functionally distinct versions of these channels in single neurons, allowing complex spatiotemporal patterning of electrical signals. For example, vertebrates use distinct suites of voltage-gated K^+^ channels in axons and dendrites (Trimmer, 2015). While ctenophores have large numbers of voltage-gated K^+^ channel genes, it is unknown whether ctenophores are more limited in the number of channel types they can express per cell. The key to answering this question is to determine whether ctenophore Shakers are broadly assembly compatible or whether ctenophores have independently evolved multiple assembly-incompatible subfamilies equivalent to the Kv1-4 subfamilies in cnidarians and bilaterians. Interestingly, MlShak4 and MlShak5 are not assembly compatible (Simonson et al., 2024) and thus may represent distinct subfamilies. We did not attempt to define ctenophore Shaker subfamilies boundaries here because, much like confirming silent/regulatory subunit phenotypes, this is an enormous job to address in a comprehensive way given the size of ctenophore Shaker family. Molecular dynamics modeling of T1 assembly domain interactions can be predictive of assembly compatibility in Shaker channels (Jegla et al., 2024; Simonson et al., 2024) and could aid in subfamily prediction. Future studies to address this question will provide key insights into the neuronal signaling potential of ctenophores.

## Acknowledgements

This research was supported by the Huck Institutes of the Life Sciences and the Eberly College of Science at Penn State University and an Allen Distinguished Investigator Award (to JFR), a Paul G. Allen Frontiers Group advised grant of the Paul G. Allen Family Foundation.

